# Type 2 diabetes susceptibility gene GRK5 regulates physiological pancreatic β-cell proliferation via phosphorylation of HDAC5 in mice

**DOI:** 10.1101/2022.12.23.521810

**Authors:** Shugo Sasaki, Cuilan Nian, Eric E. Xu, Daniel J. Pasula, Helena Winata, Sanya Grover, Dan S. Luciani, Francis C. Lynn

## Abstract

**Objective:** Diabetes onset is accompanied with β-cell deficiency, and thus restoring functional β-cell mass is a promising therapy for those with diabetes. However, the regulatory mechanisms controlling β-cell mass are not fully understood. Previously, we demonstrated that the transcription factor SOX4 is required for β-cell proliferation in the prediabetic state. To elucidate potential mechanisms by which SOX4 regulates β-cell mass, we performed RNA sequencing (RNA-seq) using pancreatic β-cell specific SOX4 knockout mice (βSOX4 KO). The RNA-seq revealed decreased expression of GRK5, a known type 2 diabetes susceptibility gene whose association with diabetes has not been fully elucidated. Therefore, we aimed to clarify the function of GRK5 in pancreatic β cells.

**Methods:** We generated a novel pancreatic β cell-specific GRK5 knockout mass (βGRK5 KO) and evaluated glucose tolerance and metabolic changes by body weight measurement, oral glucose tolerance test, and insulin tolerance test. The number of pancreatic β cells was quantified by immunohistochemistry. Glucose loading and Ca^2+^ imaging was performed on isolated islets to evaluate insulin secretory capacity. To elucidate the mechanism of βGRK5 on β cell mass regulation, we performed RNA-seq of isolated islets and identified the signaling pathways that could be regulated by GRK5. Furthermore, *in vitro* experiments were conducted using human islets and mouse βGRK5 KO islets to clarify the direct effects of GRK5 on these pathways.

**Results:** βGRK5 KO islets showed impaired glucose tolerance and insulin secretion, but no change in body weight or insulin resistance, suggesting that the main cause of impaired glucose tolerance is impaired insulin secretion. Isolated islets showed no abnormalities in insulin secretory capacity or changes in calcium influx, but histologically showed a decrease in β cell mass. Consistent with the decreased β cell mass in βGRK5 KO, the cell cycle inhibitor gene *Cdkn1a* was upregulated in βGRK5 KO islets; this phenocopies the βSOX4 KO. Furthermore, we found that Grk5 positively regulates facultative increases in β cell mass through activity-dependent phosphorylation of HDAC5 and subsequent transcription of immediate early genes (IEGs) such as *Nr4a1*, *Fosb*, *Junb*, *Arc*, *Egr1* and *Srf*.

**Conclusions:** Our results suggest that GRK5 is associated with type 2 diabetes through regulation of β cell mass. GRK5 could become a potential target of cell therapy to preserve functional β cells during the progression towards frank diabetes.

## 1. INTRODUCTION

Loss of functional insulin-producing β cells in the pancreas is one of the major pathologies of diabetes, and thus restoring functional β cell mass is a promising therapy for those with diabetes. However, the mechanisms of β cell mass regulation are not fully understood. The intronic SNPs of type 2 diabetes (T2D) susceptibility gene, *CDKAL1*, are hypothesized to regulate the expression of the nearby gene, *SOX4*, not *CDKAL1* itself [1,2]. SOX4 is a DNA-binding transcription factor that regulates β-cell mass through direct binding to the promoter region of the cell cycle gene, *Cdkn1a/p21*, among other targets [3,4].

G protein-coupled receptor kinases (GRKs) act to alter G protein coupled receptor (GPCR) signaling by changing the affinity of receptors for arrestins; usually they facilitate a switch in signaling through G proteins to signaling downstream of arrestin binding [5,6]. The activity of GRK5 is regulated by cell activity and calcium influx; Ca/CaM binding both increases GRK5 kinase activity and displaces the kinase from the membrane, potentially allowing GRK5 to phosphorylate cytoplasmic and nuclear targets [7–9]. GRK5 has been identified as a T2D susceptibility gene by genome-wide association study (GWAS) [10–14]. A variant of GRK5 is associated with the therapeutic efficacy of insulin secretagogue, repaglinide, in those with T2D [15], and with plasma LDL-cholesterol levels [16]. Germline *Grk5* knockout mice displayed mildly impaired glucose tolerance [17]. Despite these observations, the role of GRK5 in β cells has not been well studied.

Here, we report that β cell mass regulation by Sox4 is accompanied by a change of *Grk5* expression in islets. In order to determine the role of Grk5 *in vivo*, β cell specific *Grk5* knockout mice (βGRK5 KO) were generated and found to have impaired glucose tolerance with reduced β cell mass. In agreement with this phenotype, cell cycle inhibitor gene, *Cdkn1a* was upregulated in βGRK5 KO islets. As one potential mechanism linking kinase function to β cell proliferation, we found that Grk5 may regulate β cell mass through phosphorylation of HDAC5 and subsequent de-repression of immediate early genes (IEGs) that are important for cell growth.

## 2. RESULTS

### 2.1. Grk5 is downregulated in β cell specific Sox4 knockout islets

Sox4 ablation in β cells led to reduced β cell mass during prediabetes in mice [3]. To investigate downstream mechanism of Sox4 in the adult mouse islets, we performed RNA-sequencing (RNA-seq) using the islets of β cell specific Sox4 knockouts (βSOX4 KO). Pathway analysis showed that G-protein signaling was altered, and among the genes listed in that pathway, G protein-coupled receptor kinase 5 (Grk5) gene expression was downregulated in Sox4 KO islets (Fig. S1). GRK5 has been identified as T2D susceptibility gene in Asian populations by GWAS [10]. Thus, we hypothesized that SOX4 regulates β cell mass, at least in part, through regulation of GRK5 expression and tested this hypothesis here.

### 2.2. B cell specific Grk5 knockout mice displayed impaired glucose tolerance

To investigate a role of Grk5 in the islet, βGRK5 KO were generated by crossing *Pdx1-CreER* transgenic [18] with *Grk5 floxed* mice [19]. Tamoxifen was given at 6 weeks of age to induce knockout (Fig. 1A). Grk5 mRNA was reduced by >70% and protein expression lost in 8- and 18-week-old βGRK5 KO mice (Fig. 1B, C). Metabolic measures including body weight, fasting, and fed blood glucose levels were not altered in βGRK5 KO (Fig. 1D). However, βGRK5 KO mice became glucose intolerant with aging (Fig. 1E). Females exhibited a similar phenotype to males with milder impaired glucose tolerance (Fig. S2). As such, males were selected for further studies. By 18 weeks, fasting and fed insulin secretion were reduced in βGRK5 KO mice without changes in insulin sensitivity measured by insulin tolerance test (ITT) (Fig. 1F, G). Together, these studies suggest that β cell function or β cell mass was impaired in βGRK5 KO.

**Figure 1.**
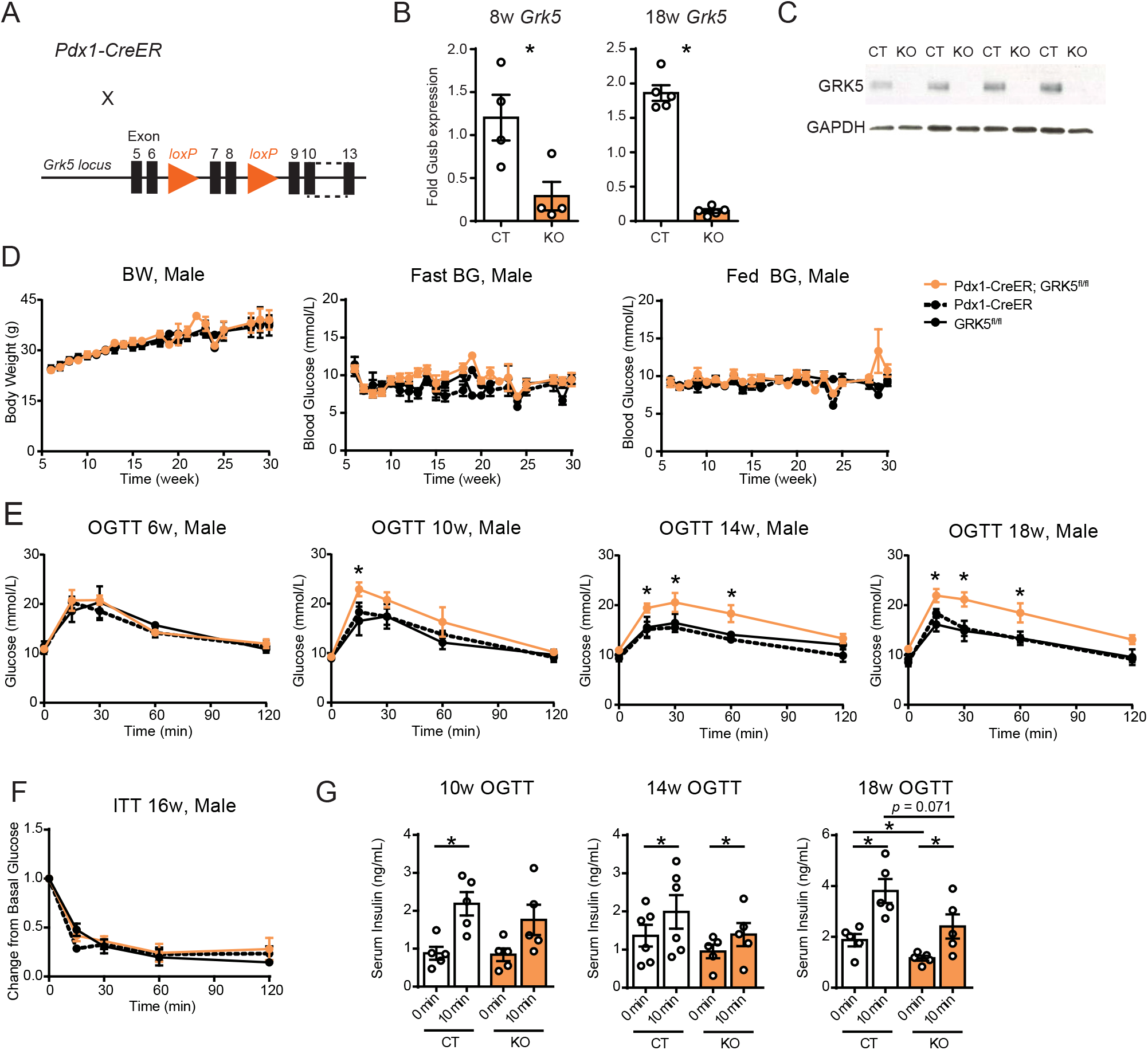
β-cell specific GRK5 knockout mice displayed impaired glucose tolerance. A) Scheme for generation of β-cell specific GRK5 knockout mice (Pdx1-CreER; GRK5^fl/fl^). \**p*<0.05. B) GRK5 mRNA expression levels in 8- and 18-week-old mice. Tamoxifen was injected at 6-week. CT: GRK5^fl/fl^, KO: Pdx1-CreER; GRK5^fl/fl^. n=4 for 8-week, n=5 for 18-week. C) Western blotting for GRK5 and GAPDH. CT: GRK5^fl/fl^, KO: Pdx1-CreER; GRK5^fl/fl^. D) Body weight, fasting blood glucose, and fed blood glucose levels over time. E) OGTT profiles for 6-, 10-, 14-, and 18-week old mice. \**p*<0.05. F) ITT profiles at 16-week. G) Serum insulin levels during OGTT. CT: GRK5^fl/fl^, KO: Pdx1-CreER; GRK5^fl/fl^. n=5-6. \**p*<0.05.

β cell function was evaluated by glucose-stimulated insulin-secretion (GSIS) using isolated βGRK5 KO mouse islets; no alterations in high glucose- or KCl-stimulated insulin secretion were observed (Fig. S3A). Total insulin in the isolated islets was unchanged (Fig. S3A). Dynamics of glucose- and KCl-regulated Ca^2+^ influx into the isolated islets were not altered (Fig. S3B). As GRKs normally phosphorylate GPCRs to reduce G protein binding and induce β-arrestin binding, we tested if knockout of Grk5 led to inappropriate activation of GLP-1 receptor by stimulating islets with GLP-1 receptor agonist Exendin-4. Neither Exendin-4-induced Ca^2+^ influx or insulin secretion was altered in the βGRK5 KO (Fig. S3A, B). Together, these studies indicate that GRK5 KO does not markedly influence β cell stimulus-secretion coupling.

### 2.3. Grk5 deletion results in reduced β cell mass with less proliferation and more apoptosis

Next, we tested whether β cell mass was affected by β cell specific Grk5 deletion. βGRK5 KO islets displayed reduced β cell numbers at 18 weeks of age (Fig. 2A, B). EdU-labelled proliferative β cells were decreased in βGRK5 KO earlier, at 8 weeks of age, 1 week following knockout (Fig. 2C, D). Apoptotic β cells assessed by TUNEL were increased in βGRK5 KO (Fig. 2E, F). To uncover mechanisms for decreased proliferation and increased apoptosis in β cells in βGRK5 KO, cell cycle-related gene expression was investigated. Expression levels of cyclin-dependent kinase inhibitor 1a gene *Cdkn1a (p21)* increased in βGRK5 KO, similar to what we previously observed in the βSOX4 KO (Fig. 2G and Fig. S1) [3]. As such, we conclude that impaired glucose tolerance in the βGRK5 KO results from reduced β cell mass driven by reduced proliferation and increased apoptosis, potentially as a result of *Cdkn1a* induction. Furthermore, the βGRK5 KO phenocopies the βSOX4 KO, suggesting that Grk5 functions downstream of Sox4 in adult β cells.

**Figure 2.**
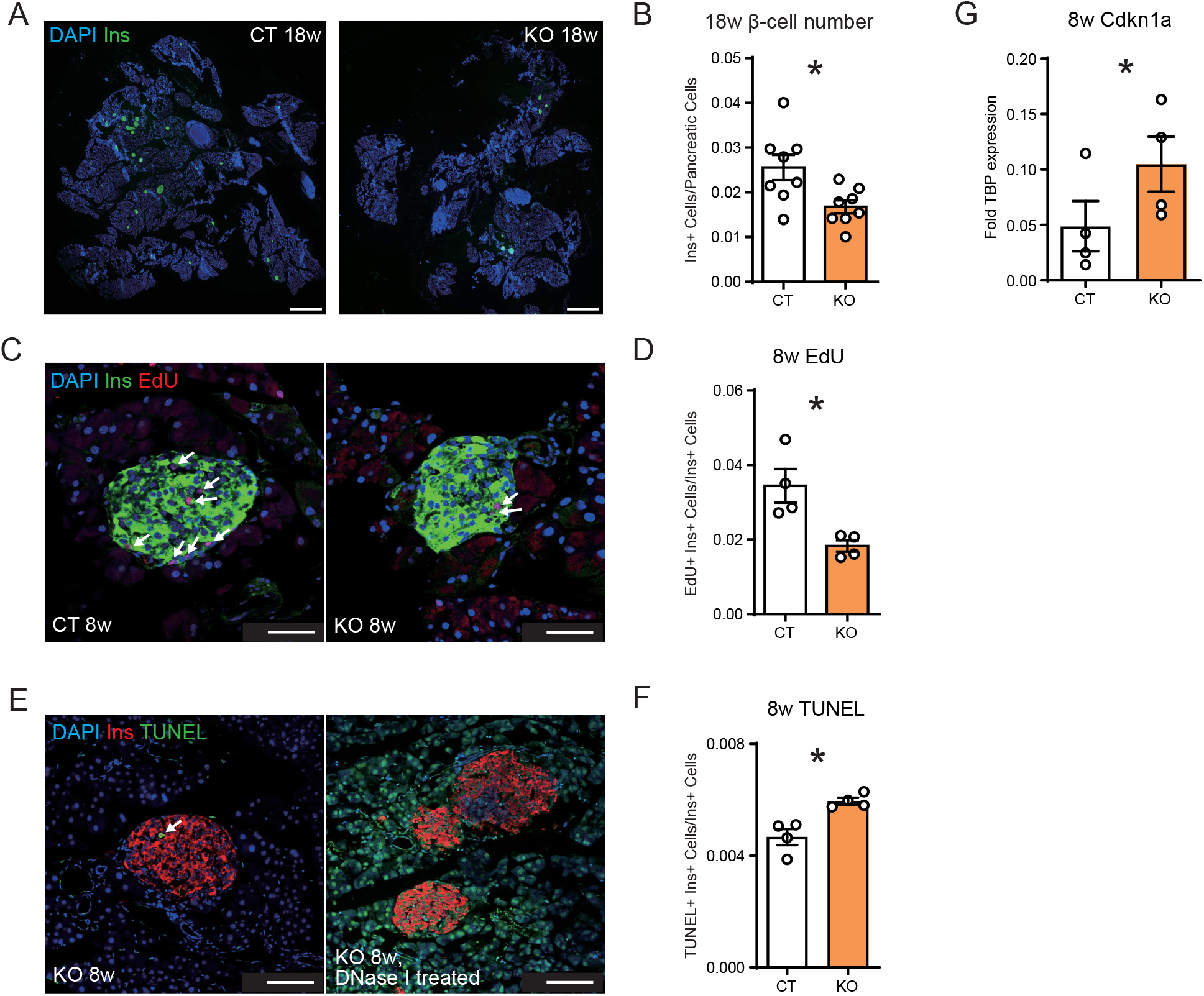
β-cell specific GRK5 knockout mice displayed reduced β-cell mass with less proliferation and more apoptosis. A) Immunohistochemical analysis for insulin in the pancreas at 18-week. Scale bar, 1 mm. B) Quantification for β-cell number. n=7. C) Immunohistochemical analysis for insulin and EdU at 8-week. Arrows indicate EdU-positive cells. Scale bar, 50 μm. D) Quantification for EdU-positive β-cell number. n=4. E) Immunohistochemical analysis for insulin and TUNEL at 8-week. Arrows indicate TUNEL-positive cells. Scale bar, 100 μm. F) Quantification for TUNEL-positive β-cell number. A DNase I treated sample is shown as a positive control. n=4. G) Cdkn1a gene expression levels in the islet. CT: GRK5^fl/fl^, KO: Pdx1-CreER; GRK5^fl/fl^. n=5-6. \**p*<0.05.

### 2.4. RNA-sequencing revealed alteration of mass regulation-related gene expressions in βGRK5 KO islets

Next RNA-sequencing (RNA-seq) was conducted to uncover the pathways downstream of GRK5 important for regulation of β cell mass. Among 17,946 detected transcripts, 601 genes were upregulated, and 398 genes were downregulated in βGRK5 KO islets (Fig. 3A). As expected, pathway analysis showed alteration of G-protein-related signaling (Fig. 3B). Of note, the most significantly changed pathway was PI3 kinase pathway, in which *Pik3r1, Pik3ca, Prkca, Gsk3b*, and *Sos2* genes were included (Fig. 3B, C). In addition to the alteration of AKT and ERK signalling pathways, IEGs such as *Arc*, *Fosb*, *Junb*, and *Nr4a1* were downregulated in both βGRK5 KO and βSOX4 KO islets, suggesting that appropriate expression levels of IEGs are necessary for β cell mass regulation in accordance with a previous reports (Fig. 3A, C) [20]. Furthermore, 15 transcripts affected by Grk5 ablation have been reported as T2D susceptibility genes, indicating their potential involvement in β cell mass regulation (Fig. 3 and S4).

**Figure 3.**
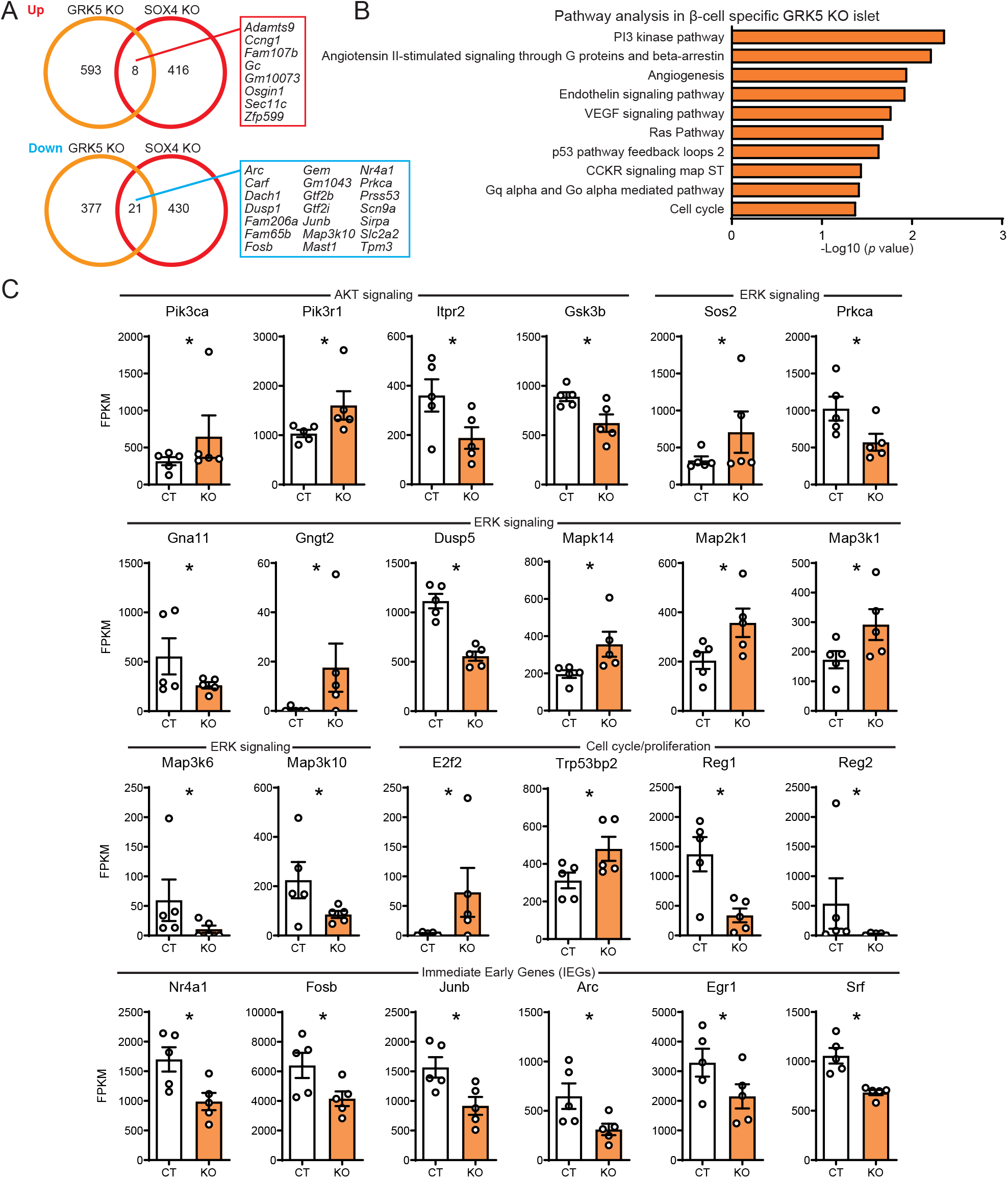
RNA-sequencing revealed alteration of mass regulation-related gene expressions in β-cell specific GRK5 knockout mice. A) The number of differentially expressed genes (DEGs) in GRK5 and SOX4 knockout islets. Overlapped DEGs are shown. B) Panther pathway analysis in β-cell specific GRK5 KO islets at 18-week. Pathways with *p*<0.01 are shown. C) FPKM for DEGs related to AKT signaling, ERK signaling, cell cycle/proliferation, and IEGs are shown. CT: GRK5^fl/fl^, KO: Pdx1-CreER; GRK5^fl/fl^. n=5. \**p*<0.05.

### 2.5. Phosphorylation of HDAC5, but not ERK and AKT signaling, is altered in βGRK5 KO

In order to test whether Grk5 modulates GPCR signaling in β cells, we investigated the AKT and ERK pathways, in response to activation of the β cell GLP-1R. Grk5 knockout did not significantly change AKT or ERK phosphorylation by multiple GLP-1R agonists that have previously been shown to signal predominantly through the PI3K/AKT (GLP-1, exendin-4) or the β-arrestin/ERK (exendin-4, oxyntomodulin) pathways [21–23]. Notably, phospho-ERK trended towards being elevated in the Grk5 KO (Fig. S5) suggesting that GRK5 may indeed be important for regulating β arrestin binding to activated GLP-1 receptors. However, as ERK signaling is thought to drive increases in β cell mass and the βGRK5 KO islets had reduced mass it seems unlikely that GRK5 modulation of GLP-1R signaling contributes to beta cell proliferation [24].

GRK5 functions by phosphorylation of G-proteins in order to regulate β-arrestin recruitment and ERK signaling but also phosphorylates non-canonical pathways. For example, GRK5 phosphorylation of HDAC5 leads to de-repression of MEF2 transcriptional activity, and cellular proliferation in cardiomyocytes, neurons and other cell types through induction of immediate early genes (IEGs) [6,7,25] [26].

Among HDAC family members, HDAC5 was most highly expressed in mouse islets (Fig. S6A). Thus, we investigated whether GRK5-HDAC5-MEF2-IEG axis is active in β cells. First, βGRK5 KO islets displayed less phosphorylation of HDAC5 (Fig. 4A). Secondly, GRK5-driven phosphorylation of HDAC5, which is induced acutely by glucose and KCl-stimulation, was suppressed in βGRK5 KO islets (Fig. 4B). Thirdly, in human islets, stimulation-induced phosphorylation of HDAC5 was blocked by inhibiting Grk5 with amlexanox (Fig. 4C) [27]. In sum these data, along with the observed downregulation of various previously characterized HDAC5 target IEGs revealed by the RNA-seq (Fig. 3C), suggest that GRK5 positively regulates β cell mass expansion through phosphorylation of HDAC5.

**Figure 4.**
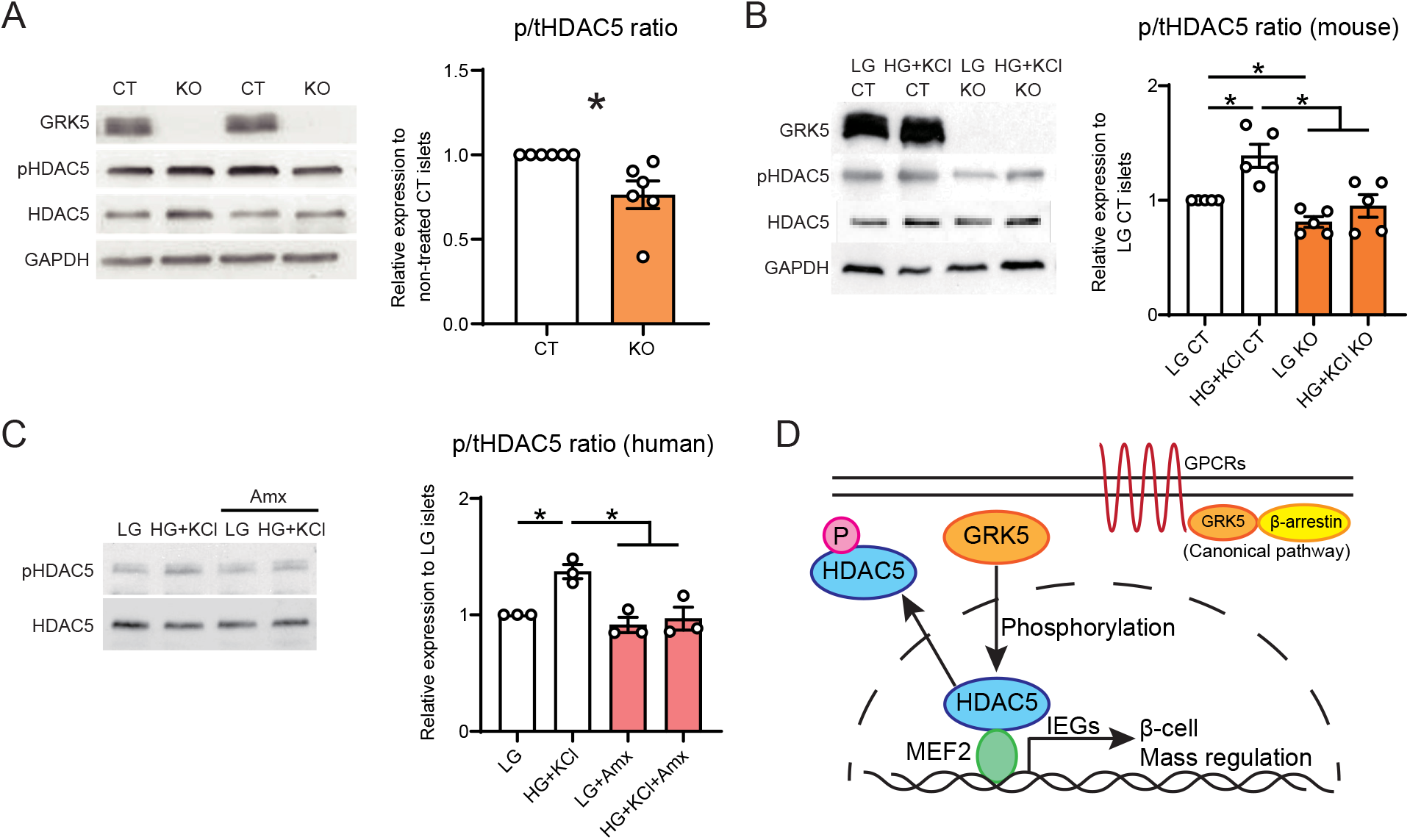
Phosphorylation of HDAC5 is altered in β-cell specific GRK5 knockout mice and human islets treated with amlexanox. A) Western blotting for HDAC5 phosphorylation. 80 mouse islets (18-week) were applied to each lane. CT: GRK5^fl/fl^, KO: Pdx1-CreER; GRK5^fl/fl^. p/tHDAC5 ratio: phospho HDAC5/total HDAC5 ratio. n=6. \**p*<0.05. B) Western blotting for HDAC5 phosphorylation status after stimulation. 80 mouse islets (18-week) were applied to each lane. The islets were treated by LG (2.8 mM low glucose) and HG + KCl (16 mM high glucose + 30 mM KCl) for 1 hr. n=5. \**p*<0.05. C) Western blotting for HDAC5 phosphorylation status after stimulation. 100 human islets were applied to each lane. The islets were treated by LG, HG+KCl and Amx (50 μM amlexanox) for 1 hr. n=4. \**p*<0.05. D) Scheme for β-cell mass regulation by GRK5. HDAC5 is suppressing activity of MEF2 that transcribes IEGs and regulates cell mass. GRK5 inactivates HDAC5 by phosphorylation, and de-suppresses MEF2 activity, resulting in β-cell mass expansion.

## 3. DISCUSSION

In this study, we found that GRK5 is downstream of SOX4 and is important for regulation of β cell mass in mice. To our knowledge, this is the first report demonstrating the role of the T2D susceptibility gene *GRK5* in β cells. Whereas β cell function was not altered by GRK5 KO, β cell proliferation was decreased and apoptosis were increased, along with upregulation of cell cycle arrest genes. Phosphorylation of HDAC5 by GRK5 and subsequent upregulation of IEGs were revealed as a potential mechanism of β cell mass regulation. Further, we found that GRK5 is necessary for phosphorylation of HDAC5, which is induced by acute glucose and KCl-stimulation, in the mouse and human β cell.

*GRK5* has been identified as T2D susceptibility gene in the Chinese population [10]. In general, East Asians show a higher insulin sensitivity and a lower acute insulin response than other races [28], which could be partially explained by reductions in GRK5 function and reduced β cell mass, as shown in mice here. Another study revealed the higher frequency of the GRK5 rs10886471 T allele polymorphism in Chinese with T2D than in healthy subjects. Furthermore, patients with the TT genotype (at rs10886471) were resistant to treatment by insulin secretagogue, repaglinide (a lower reduction of fasting plasma glucose and lower differential values of postprandial serum insulin) compared with those with CC and CT genotypes [15]. This suggests that the key role GRK5 plays in regulating β cell mass and/or function is conserved between humans and mice.

GRK5 has been actively investigated in chronic degenerative diseases such as heart failure, neurological disorders, and cancer [6,29]. Notably, GRK5 promotes cell proliferation pathologically in cardiac hypertrophy [7,8,25]. Additionally, GRK5 catalytic activity is essential to preserve cardiomyocyte survival by preventing p53-induced apoptosis [30]. Notably an *in vitro* study demonstrated that knockdown of GRK5 resulted in increased p53 and CDKN1A expression, and showed that GRK5 controlled microtubule nucleation and normal cell cycle progression [31]. Taken together with our data (*e.g*. Fig. 3), it appears that GRK5 regulates cell cycle physiologically and pathologically in tissue- and context-specific manner. Currently, the development of small-molecule inhibitors targeting GRK5 for the treatment of heart failure and other conditions has attracted wide attention [29], however, it is possible that GRK5 inhibition may have long-term unfavorable effects on β cell mass and diabetes risk.

More than 800 GPCRs have been identified in rodents and humans while only seven GRKs work to tune GPCR signaling. Expression analyses show that of the GRK-family transcripts, GRK4-6 are most abundantly expressed GRKs in both mouse and human islets (Fig. S6B, C). As GLP-1R signaling is important for normal β-cell function, we tested whether Grk5 loss altered the AKT or ERK pathways and found no significant effects. Other GPCRs such as the M3R and GPR40 could be regulated by Grk5 in the β cell, and we will be interested in testing this in future studies [32]. In addition to HDAC5, Grk5 may phosphorylate other immediate early gene targets in β cells. IEGs regulate cell cycle genes and cell mass in various tissues [26]. It is controversial whether Nr4a1 alone regulates β cell mass [33,34]; however, a set of IEGs including *Srf*, *Junb*, *Fos*, and *Egr1* have been shown to induce β cell proliferation [20]. The role of IEGs in β cell proliferation may change depending on the pathophysiological context and orchestrated control of IEGs through manipulation of GRK5 activity may be another target for diabetes therapy.

In conclusion, we discovered that T2D susceptibility gene *GRK5* regulates physiological pancreatic β cell proliferation, possibly through phosphorylation of HDAC5 and transcription of IEGs. Future studies will aim to understand whether GRK5 is a tractable β cell drug target.

## 4. METHODS

### 4.1. Animals and human islet studies

All experiments were approved by the University of British Columbia Animal Care Committee. Animals were housed under a 12-h light/dark cycle, fed ad libitum with standard chow diet (5010; Lab Diets). Mouse strains used were on the C57BL/6 background. Pdx1-CreER; Sox4^flox/flox^ mice were generated as described [3,18]. Grk5^flox/flox^ mice were generated by crossing FLP mice [35] with Grk5^tm1a(KOMP)Mbp^ mice [19] obtained from International Mouse Phenotyping Consortium. Grk5^flox/flox^ mice were crossed with Pdx1-CreER mice [18] to generate Pdx1-CreER; Grk5^flox/flox^ experimental mice. To control for potential tamoxifen effects on β cell replication, all mice were administered 8 mg tamoxifen in corn oil (60 mg/mL) by oral gavage every other day beginning at 6 weeks of age. Human islet studies were approved by the BC Children’s and Women’s Hospital Research Ethics Board. Islets were provided by the University of Alberta Islet Distribution Program. Human islet data is provided in the attachment.

### 4.2. Metabolic phenotype analysis

For glucose tolerance tests, mice were weighed and fasting saphenous vein blood glucose levels obtained using a OneTouch UltraMini glucometer (Lifescan) after a 10-h fast during the dark cycle. 2 g/kg D-glucose was delivered via intraperitoneal (i.p.) injection and blood glucose levels determined at 0, 15, 30, 60, and 120 min. Blood was collected at fasting and 10 min following i.p. glucose for determination of serum insulin levels using ELISA (STELLUX; ALPCO Salem, NH). Blood glucose levels greater than the glucometer detection limit were reported as 33.3 mmol. Intraperitoneal insulin tolerance test was performed following a 3-h fast during the light cycle with 1 unit/kg i.p. insulin injection.

### 4.3. Immunohistochemical analysis

Immunostaining was performed on 5-μm paraffin sections as described [3]. Primary antibodies were applied overnight at 4°C in PBS-0.3% Triton+ 5% horse serum. After washing with PBS, secondary antibodies and DAPI were applied for 1h at room temperature, before mounting on slides using SlowFade Diamond Antifade (Invitrogen). A primary antibody was guinea pig anti-insulin (1:1000; A0564; Dako). Secondary antibodies were donkey anti-guinea pig FITC and Cy3 (1:200; Jackson Immuno Research Laboratories, #706-096-148 and # 706-166-148). 5-Ethynyl-2’-deoxyuridine (EdU) staining on sections was performed as described [3]. A total of 0.5 mg EdU was administered (i.p.) twice a day for 7 days, beginning at 7 weeks of age. TdT-mediated dUTP-X nick end labeling (TUNEL) was performed using In Situ Cell Death Detection Kit, TMR red (Roche). Twelve sections (250-μm intervals) of each pancreas were imaged using a BX61 microscope and tiled using the cellSens Dimension software (Olympus). CellProfiler 3.0 [36] was used to quantify images, and counts were normalized to total pancreatic nuclei. For EdU and TUNEL quantification, sections were imaged using confocal microscopy (Leica SP8; Leica Microsystems) and counted as above.

### 4.4. Ex vivo islet assays

Mouse pancreatic islets were isolated and recovered overnight. Human islets were recovered overnight after receipt. Following preincubation in Krebs-Ringer buffer containing 2.8 mM glucose for 1 h at 37°C, 100 islets were incubated in either 2.8, 25 (±50 nM exendin-4), or 2.8 mM glucose with 40 mM KCl for 1 h at 37°C. Supernatants were collected for insulin measurement. Fura-2 calcium imaging was performed as described previously [37].

### 4.5. Gene expression assays

RNA was isolated with TRIzol, DNase treated (Turbo DNAse Free; Thermo Fisher Scientific), and reverse transcribed with Superscript III (Thermo Fisher Scientific) as previously described [3]. TaqMan quantitative PCR (qPCR) was performed as described [3]. Primers used were: mouse Grk5 (forward [F], 5′-GCACTCAACGAAAAGCAGATTC-3′; reverse [R], 5′-GTGCATCTTTGGTTTCATAGGC-3′; and probe 5′-AGGTCAACAGCCAGTTTGTGGTCA-3′), mouse Cdkn1a (F, 5′-CTGAGCGGCCTGAAGATT-3′; R, 5′-ATCTGCGCTTGGAGTGATAG-3′; and probe 5′-AAATCTGTCAGGCTGGTCTGCCTC-3′), mouse Gusb (F, 5′-TCTAGCTGGAAATGTTCACTGCCCTG-3′; R, 5′-CACCCCTACCACTTACATCG-3′; and probe 5′-ACTTTGCCACCCTCATCC-3′), human GRK5 (F, 5′-AGGTCTTCACACTGGCTAATG-3′; R, 5′-CAATGGAGCTGGAAAACATCG-3′; and probe 5′-AGGAAAGCGCAAAGGGAAAAGCAAG-3′), and human TBP (F, 5′-TGGGATTATATTCGGCGTTTCGGGC-3′; R, 5′-GAGAGTTCTGGGATTGTACCG-3′; and probe 5′-ATCCTCATGATTACCGCAGC-3′).

### 4.6. RNA-sequencing

Following total RNA extraction and DNase treatment, samples were rRNA depleted utilizing the RiboGone Mammalian-Low Input Ribosomal RNA Removal Kit (Takara Bio USA Inc., Mountain View, CA, USA), and Agencourt AMPure XP SPRI beads (Beckman Coulter). rRNA depleted total RNA was assessed for rRNA depletion, RNA integrity, size distribution, and concentration prior to library generation with the Agilent RNA Pico 6000 kit on the Agilent 2100 Bioanalyzer (Agilent Technologies, Santa Clara, CA, USA). rRNA depleted RNA passing quality control was used in the SMARTer Stranded RNA-Seq Kit (Takara Bio USA). Following RNA incubation with stranded N6 primer, buffer, and water, samples were incubated at 94°C for 4 minutes, appropriate for samples with RIN values between 4-7. Following purification of first-strand cDNA, libraries were PCR amplified in accordance with their input RNA amount and purified. Final adapter-ligated cDNA was quantitated by Qubit 3.0 dsDNA high-sensitivity assay kit (Thermo Fisher Scientific) together with qPCR amplification using standard curves of a known concentration of adapter-ligated libraries. Sequencing was performed on the NextSeq 500 with the High Output Reagent Cartridge v2 150 cycles (75bp x 2) (Illumina, San Diego, CA, USA). Linux Ubuntu operating system version 14.04 LTS was used for analyses. Fastq files from each sample were concatenated. ENSEMBL Mus musculus genome GRCm38p.4 fastq and gtf annotation file were used for alignments. STAR v2.4.0.1 230 was utilized to index the genome fastq files, align sample transcriptomes against the genome, and output as sorted by coordinate BAM files. Counts were generated from BAM files with HTSeq [38] against annotated genes from GRCm38p.4. Finally, differential gene analysis was performed between control and experimental samples using DESeq2 [39], with an FPKM criteria of ≥5 in two or more samples, applying a Wald test on *p*-values from genes passing filtering, and adjusted for multiple testing via Benjamini and Hochberg procedure. Panther pathway analysis performed for differentially expressed genes [40].

### 4.7. Western blot

Western blots were performed as described [41]. Primary antibodies used included: mouse anti-GRK5 (D-9, 1:1,000; SCBT, sc-518005), mouse anti-phospho Erk1/2 (E10, 1:1,000; CST, 9106S), mouse anti-Erk1/2 (3A7, 1:1,000; CST, 9107S), rabbit anti-phospho Akt (1:1,000; CST, 9271S), rabbit anti-Akt (1:1,000; CST, 9272S), rabbit anti-phospho HDAC5 (1:1,000; Abcam, ab47283), mouse anti-HDAC5 (B-11; 1:1,000, SCBT, sc-133106), and mouse anti-GAPDH (1;100,000; Sigma-Aldrich, G8795). Secondary horseradish peroxidase– conjugated antibodies were from Jackson ImmunoResearch Laboratories (#115-035-174 and #111-035-008). ImageJ (NIH) was used for quantification.

### 4.8. Data sources

Type 2 diabetes susceptibility genes are listed using the NHGRI-EBI GWAS Catalog [12]. GRK1-7 gene expression data in human islets and β cells are obtained from the GEO database (accession GSE73433 [42], GSE140403 [43], and GSE57973 [44]).

### 4.9. Statistical analyses

Measurements were performed on independent samples unless otherwise specified. Statistical analyses were performed using the Prism 9.0 (GraphPad Software, La Jolla, CA). Multiple groups were analyzed by one-way ANOVA with a multiple comparison test and the Tukey-Kramer’s post-hoc test was used to compare different groups. A *p*-value < 0.05 was considered to indicate a statistically significant difference between two groups. Data are presented as the mean ± SEM.

## CREDIT AUTHOR CONTRIBUTION

S.S., C.N., E.E.X., D.J.P., H.W., and S.G. generated and analyzed data. S.S., D.S.L., and F.C.L. designed experiments. S.S. and F.C.L. drafted the manuscript. All the authors approved the version of the manuscript to be published. F.C.L. is the guarantor of this work and, as such, had full access to all the data in the study and takes responsibility for the integrity of the data and the accuracy of the data analysis.

## DATA AVAILABILITY

Data was made publicly available on GEO (accession number: GSE___).

## ACKNOWLEDGMENT

The authors thank the members of the Lynn Laboratory (Vancouver, British Columbia, Canada) for technical support, discussion, and critical reading of the manuscript. F.C.L. was supported by the Canadian Institutes of Health Research (PJT-156377) and a JDRF Career Development Award. Salary (F.C.L.) was supported by the Michael Smith Foundation for Health Research (#5238 BIOM) and the BC Children’s Hospital Research Institute. Fellowship support was provided by the Juvenile Diabetes Research Foundation (S.S.; 3-PDF-2018-587-A-N), the Michael Smith Foundation for Health Research (S.S.; 17045), and the Manpei Suzuki Diabetes Foundation (S.S.).

The authors acknowledge that UBC and BC Children’s Hospital are situated on the traditional, ancestral and unceded territories of the Coast Salish peoples – the Sḵwx̱ wú7mesh (Squamish), Səl̓ ílwətaʔ/Selilwitulh (Tsleil-Waututh) and xwməθkwəy̓ əm (Musqueam) Nations.

## APPENDIX A. SUPPLEMENTARY DATA

Supplementary data to this article can be found online at ___

---

**Figure S1.**
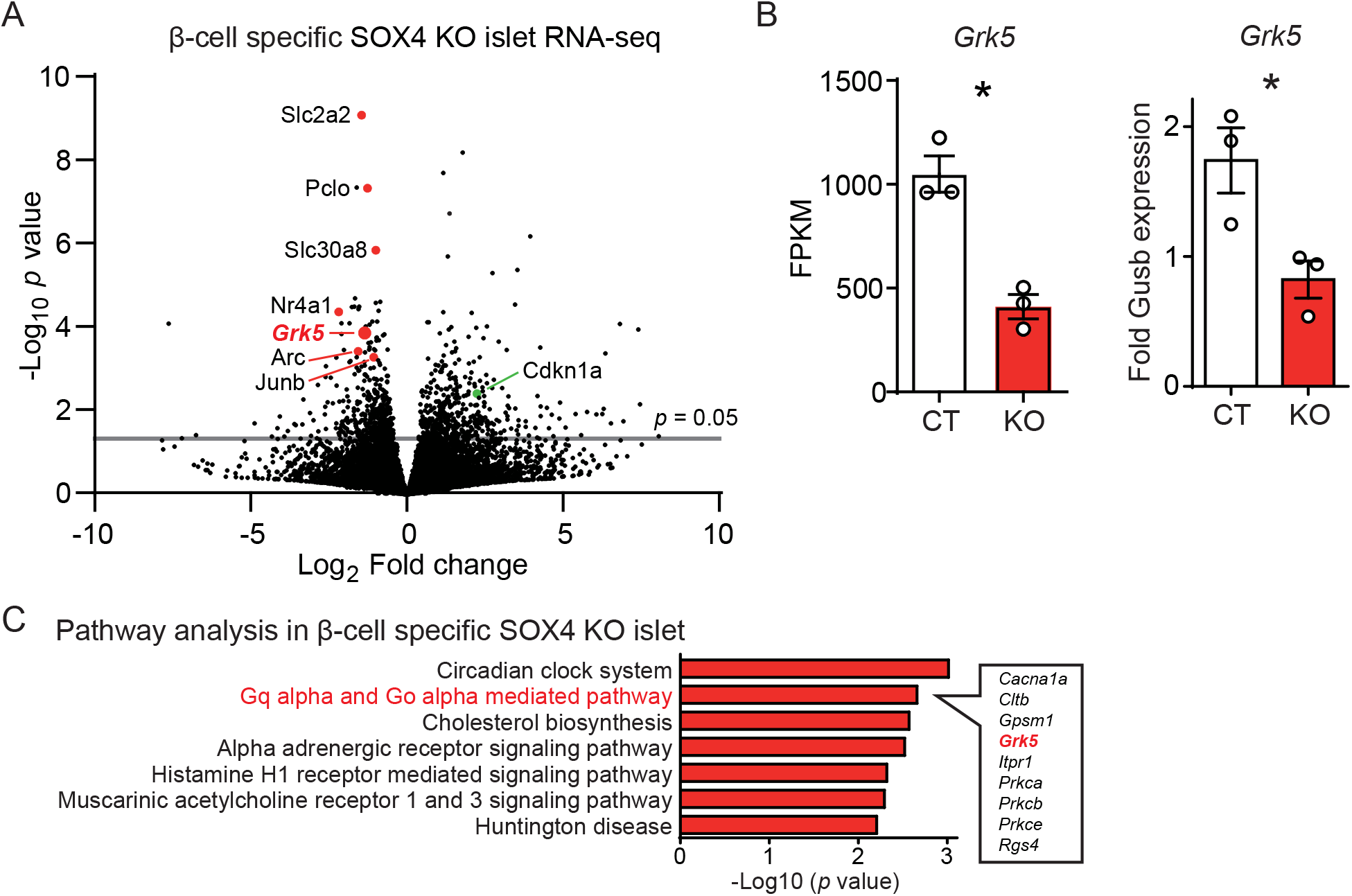
Grk5 is downregulated in β-cell specific SOX4 knockout islets A) Volcano plots for gene expressions in β-cell specific SOX4 KO islets at 8-week. B) GRK5 gene expression levels in β-cell specific SOX4 KO islets analyzed by RNA-seq and Taqman qPCR. n=3. * *p*<0.05. CT: SOX4^fl/fl^, KO: Pdx1-CreER; SOX4^fl/fl^. C) Panther pathway analysis in β-cell specific SOX4 KO islets at 8-week. Pathways with *p*<0.01 are shown. Genes in “Gq alpha and Go alpha mediated pathway” category are shown.

**Figure S2.**
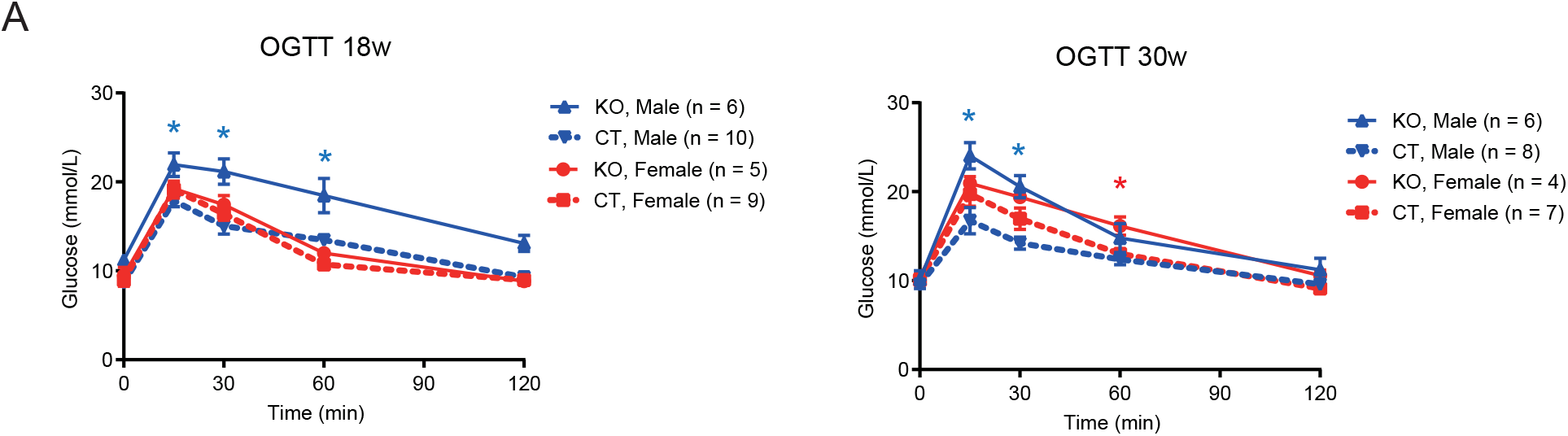
Mild impairment of glucose tolerance in female β-cell specific GRK5 knockout mice A) OGTT profiles for 18- and 30-week-old mice. CT: GRK5^fl/fl^, KO: Pdx1-CreER; GRK5^fl/fl^. \**p*<0.05.

**Figure S3.**
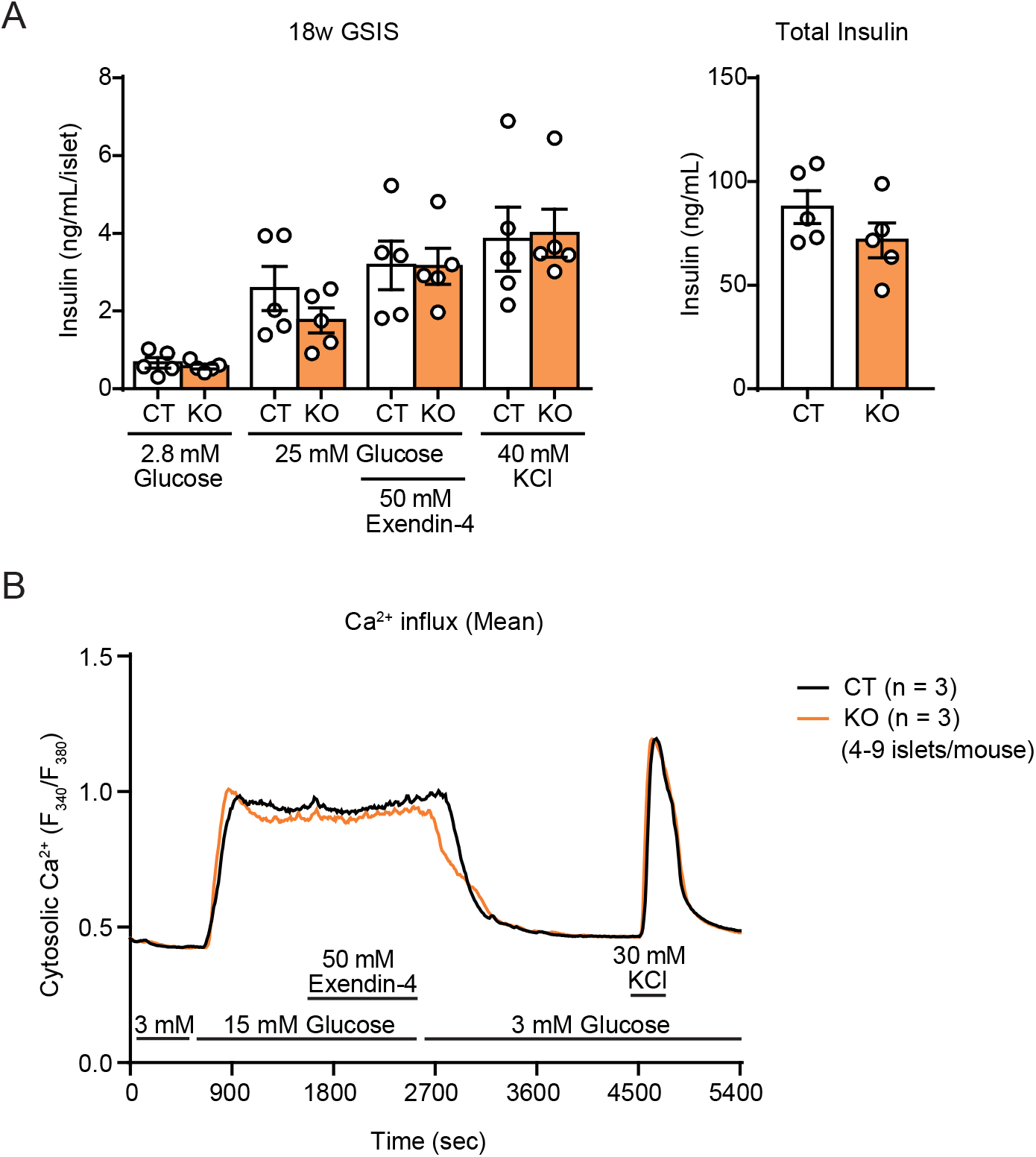
β-cell function in β-cell specific GRK5 knockout mice was not altered A) Glucose-stimulated insulin secretion and total insulin concentration with isolated islets. 100 islets were used for each. n=5. B) Mean values of cytosolic Ca^2+^ concentration with isolated islets. CT: GRK5^fl/fl^, KO: Pdx1-CreER; GRK5^fl/fl^.

**Figure S4.**
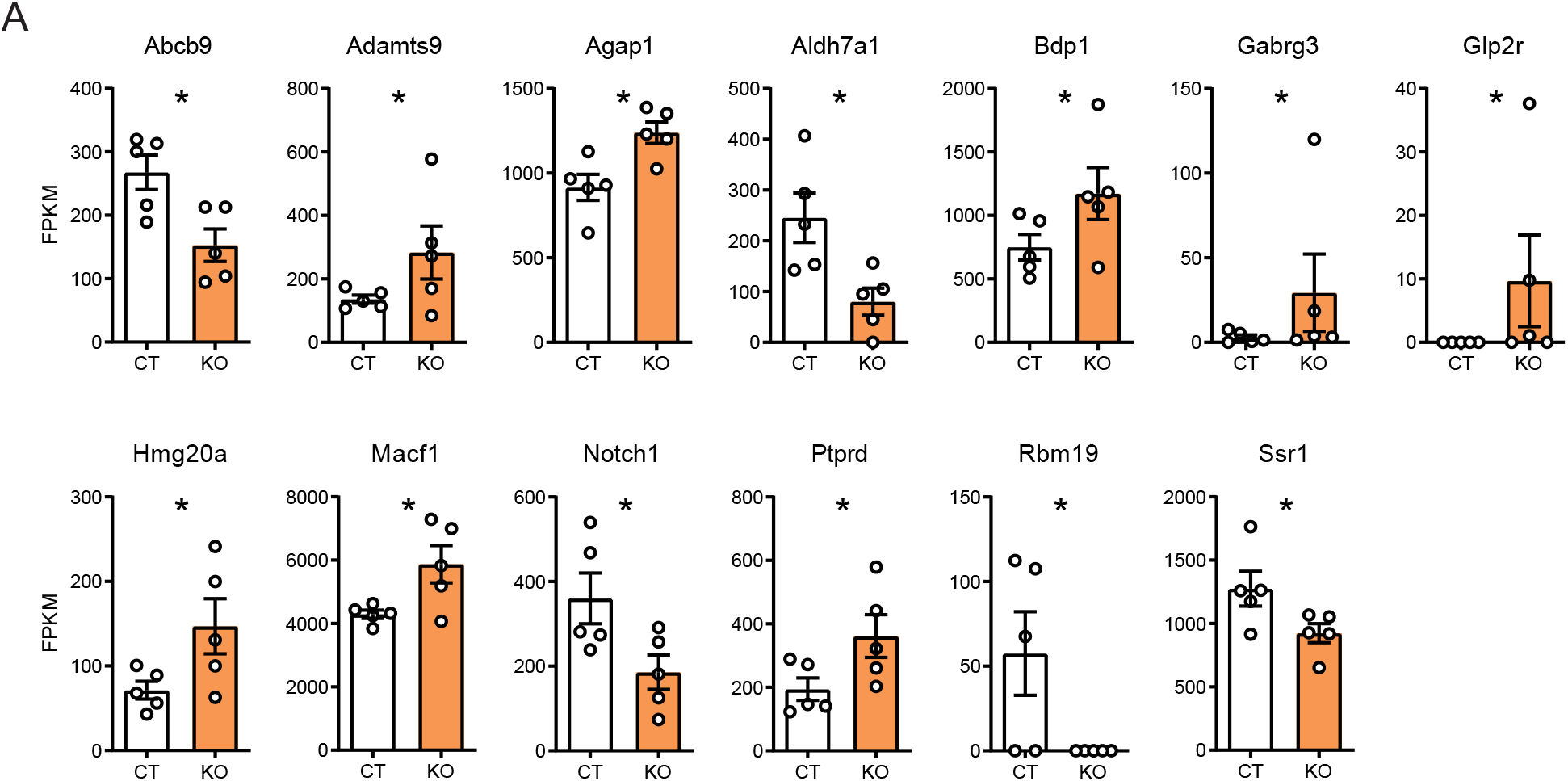
RNA-sequencing revealed alteration of type 2 diabetes susceptibility gene expressions in β-cell specific GRK5 knockout mice A) Differentially expressed genes (DEGs) by RNA-seq that are reported as type 2 diabetes (T2D) susceptibility genes by GWAS studies are shown. Mapk14 and Map3k1, that are also differentially expressed and reported as T2D susceptibility genes, are displayed in Figure 3. CT: GRK5^fl/fl^, KO: Pdx1-CreER; GRK5^fl/fl^. n=5. \**p*<0.05.

**Figure S5.**
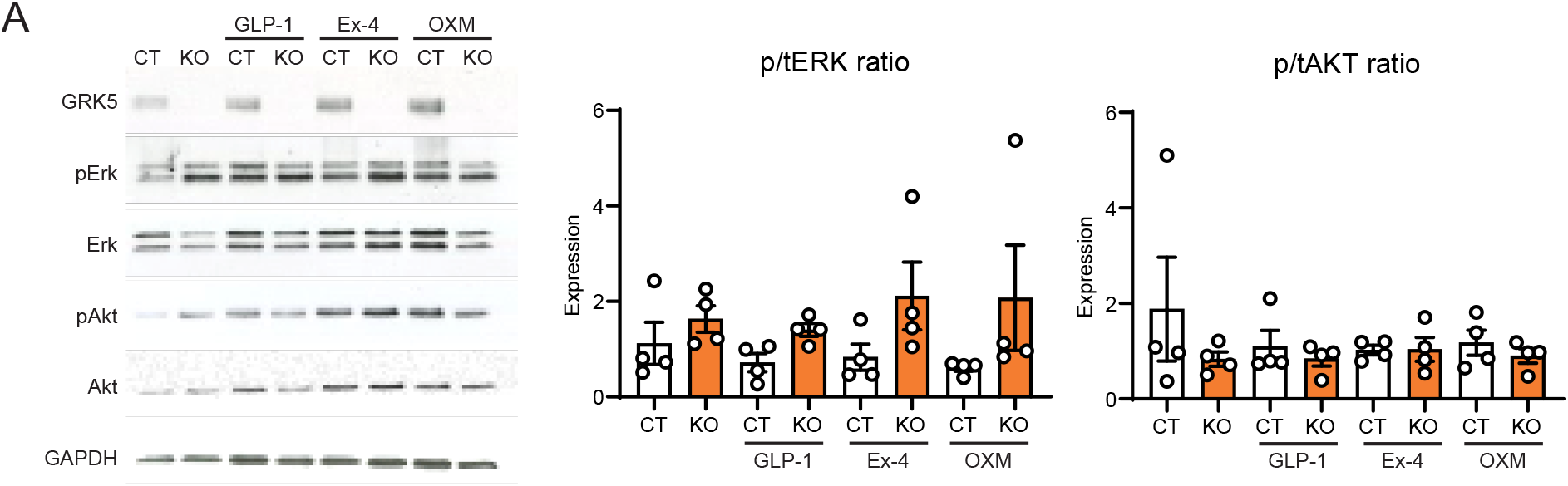
Phosphorylation of ERK and AKT is not altered in β-cell Specific GRK5 Knockout Mice. A) Western blotting for ERK and AKT phosphorylation. 80 mouse islets (18-week) were applied to each lane. The islets were treated by 100 nM of GLP-1, exendin-4 (Ex-4), and oxyntomodulin (OXM) for 1 hr.

**Figure S6.**
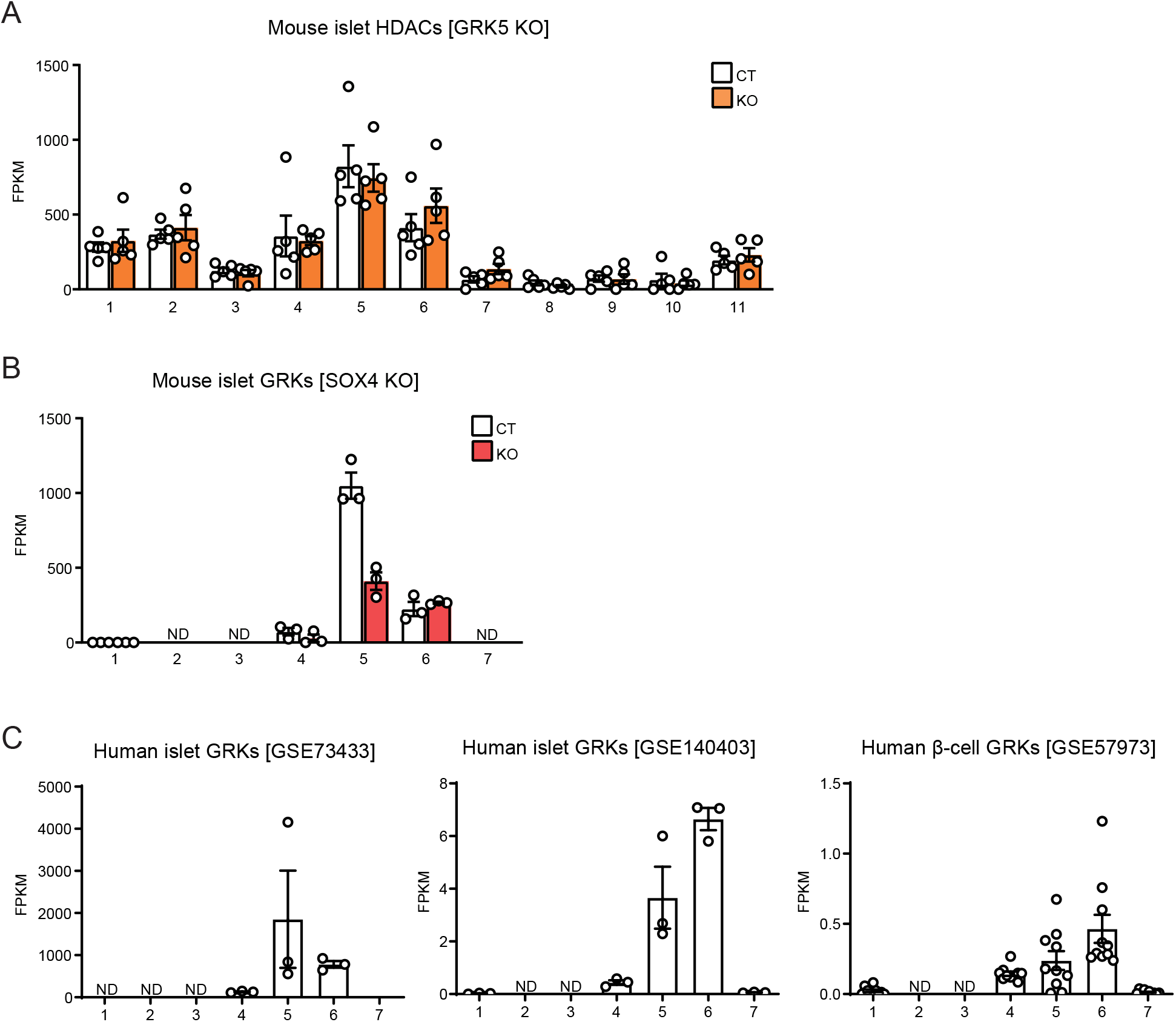
HDAC5 was mainly expressed in mouse islets, and GRK4-6 were in mouse and human islets. A) FPKM of HDAC1-11 in mouse islets. CT: GRK5^fl/fl^, KO: Pdx1-CreER; GRK5^fl/fl^. n=5. B) FPKM of GRK1-7 in mouse islets. [SOX4KO] in red. CT: SOX4^fl/fl^, KO: Pdx1-CreER; SOX4^fl/fl^. n=3. C) FPKM of GRK1-7 in human islets and β cells extracted from three accessible datasets.

